# Analysis of stress responses in *Astyanax* larvae reveals heterogeneity among different populations

**DOI:** 10.1101/2020.04.06.027649

**Authors:** Jacqueline SR Chin, Cody L. Loomis, Lydia T. Albert, Shirley Medina-Trenche, Johanna Kowalko, Alex C. Keene, Erik R. Duboué

## Abstract

Stress responses are conserved physiological and behavioral outcomes as a result of facing potentially harmful stimuli, yet in pathological states, stress becomes debilitating. Stress responses vary considerably throughout the animal kingdom, but how these responses are shaped evolutionarily is unknown. The Mexican cavefish has emerged as a powerful system for examining genetic principles underlying behavioral evolution. Here, we demonstrate that cave *Astyanax* have reduced behavioral and physiological measures of stress when examined at larval stages. We also find increased expression of the glucocorticoid receptor, a repressible element of the neuroendocrine stress pathway. Additionally, we examine stress in three different cave populations, and find that some, but not all, show reduced stress measures. Together, these results reveal a mechanistic system by which cave-dwelling fish reduced stress, presumably to compensate for a predator poor environment.

**Research Highlight:**

- Cavefish populations of *A. mexicanus* have reduced stress relative to surface conspecific at larval stages
- We show that a glucocorticoid receptor, a negative regulator of the neuroendocrine stress axis, is upregulated in stress-resistant cavefish
- There exists much ontological heterogeneity between different cavefish populations.

## Introduction

Many physiological and behavioral responses to stressful stimuli are evolutionarily conserved throughout the animal kingdom, and function to increase alertness and promote survival in times of adversity (Campos et al., 2013; Cockrem, 2013). Among diverse species, acute stress can be defined by changes in behavior, including reduced exploration and freezing, as well as an increase in circulating levels of catecholamines and glucocorticoids, the primary stress hormones found among vertebrates (Collier et al., 2017).

While behavioral and physiological responses to stress are widely thought to be adaptive (Lacey and Lacey, 1958; Schneiderman et al., 2005), pathological stress becomes debilitating and can result in various stress and anxiety related disorders (Hadany et al., 2006). Moreover, it is not clear what predisposes some individuals to stress and anxiety related disorders, though epidemiological studies point to a complex relationship between genetic and environmental components (Etkin and Wager, 2007; Koenen et al., 2008; Cornelis and Nugent, 2010). Despite these findings, the environmental factors that shape stress and how naturally occurring genetic variation contributes to physiology and the susceptibility of related diseases states remains poorly understood. A central impediment to addressing this question has been the lack of a model in which stress can be examined, and the responses correlated with naturally occurring genetic variation and a well-established ecological and evolutionary history.

The Mexican tetra, *Astyanax mexicanus*, is a small fresh-water fish native to northeast Mexico and regions of southern Texas, which provides a powerful system for studying behavioral evolution (Yoshizawa et al., 2010; Duboué et al., 2011; Elipot et al., 2013; Kowalko et al., 2013b; Jaggard et al., 2018). This species exists as an eyed, river-dwelling form, and at least 29 populations of cave-dwelling morphs (Mitchell et al., 1977; Borowsky, 2008a; Jeffery, 2009). Cave and surface populations are interfertile, allowing for genetic analysis of the complex multi-locus traits that distinguish the different surface- and cave-dwelling populations (Wilkens, 1971; Protas et al., 2006, 2008; Yoshizawa et al., 2015). The ecology and environmental setting of the caves and rivers has been well-described (Mitchell et al., 1977). Additionally, an exhaustive phylogenetic history of the species (Mitchell et al., 1977; Ornelas-García et al., 2008; Bradic et al., 2012, 2013), combined with the ability to compare between independently evolved populations provides the opportunity to compare evolutionarily-derived changes in behavior with ecology (Keene et al., 2015). Lastly, because cavefish are derived from wild-caught populations, these animals represent a genetic model system with intact naturally occurring genetic variation (Dowling et al., 2002; Borowsky, 2008b; Bradic et al., 2012; Gross, 2016; Rohner, 2018). In addition to the number of morphological traits that differ between the cave and surface morphs, several behaviors, including sleep, schooling, foraging, and aggression have been shown to differ significantly among epigean and subterranean forms (Patton et al., 2010; Yoshizawa et al., 2010; Duboué et al., 2011; Elipot et al., 2013; Kowalko et al., 2013b; a; Aspiras et al., 2015; Jaggard et al., 2017, 2018).

The zebrafish *Danio rerio*, and a number of other teleost species, have emerged as powerful models for addressing neuronal and genetic principles underlying the regulation of stress (Cachat et al., 2010; Maximino et al., 2010, 2014; Schreck et al., 2016; Sakamoto et al., 2017; Laberge et al., 2019). Studying stress in fish not only has implications in aquaculture and farming (Sneddon et al., 2016), the principles, neuronal circuitry, and neuroendocrine systems that modulate these behaviors are largely analogous with mammalian systems (Mueller et al., 2011; Biran et al., 2015), and more than 80% of disease-causing genes are conserved (Varga et al., 2018). The neuroendocrine stress axis and the primary neuroendocrine stress hormone, cortisol, are conserved between fish and humans (Collier et al., 2017). Standardized assays have been developed for studying stress responses in both adult and larval fish (Cachat et al., 2010; Maximino et al., 2010). For example, in response to a mild electric stimulus, fish, like mammals, undergo a characteristic stress response, which involves prolonged freezing followed by recovery to baseline (Duboué et al., 2017). These powerful high-throughput assays, combined with genetic tools for manipulating gene and neuronal function, has made these fish systems an invaluable asset to the study of stress biology.

We have demonstrated recently that stress in adult cavefish differs significantly from their surface counterparts (Chin et al., 2018). Adult cavefish have reduced behavioral measures of stress, but the response of larvae is unknown. In this study, we examine stress responses in larval stages of *A. mexicanus*. Our results reveal that diminished stress in cavefish is likely driven by an upregulation of the glucocorticoid receptor. We also show a divergence in ontogeny of stress states, where some cavefish populations have dampened behavioral measures of stress responses in both adult and larval stages, whereas other populations have dampened stress in adult forms, but not in larval stages. Together, our data reveal that upregulation of negative regulators of stress may underlie reduced stress states in *A. mexicanus* cavefish, and suggest that the cavefish system may be a powerful tool in examining how naturally occurring genetic variation leads to susceptibility of stress related disorders.

## Methods

### Animal husbandry

All animals used in this study were maintained on a dedicated recirculating aquatics facility (Aquaneering Inc., San Diego, CA) at Florida Atlantic University, as previously described (Borowsky, 2008c; Stahl et al., 2019a). All fish stocks originated from either the Borowsky laboratory (New York University) or the Jeffery laboratory (University of Maryland, College Park). Fish were housed in standard 10-37L glass tanks, and the water temperature held constant at 21 ± 1°C. The aquatics facility was equipped with overhead lights (25-40 lux), and the light cycle was set to 14:10 light:dark. Adult fish were fed twice a day at zeitgeber times ZT2 and ZT12 with a mixture of fish flakes (TetraMin) and black worms (Aquatic Foods, Fresno, CA) as previously described (Jaggard et al., 2018; Stahl et al., 2019a).

Larvae used in this study were generated by natural spawning of adult breeder lines, as described previously (Borowsky, 2008c). Briefly, on the night before spawning, a heater was placed in a clean tank containing male and female breeders, and the temperature was raised 4-5°C. Eggs were collected between 00:00 hr (midnight) and 04:00 hr and transferred to clean glass bowls (Pyrex). All debris and unfertilized eggs were removed from the glass bowls, and dirty water was replaced with clean water. Eggs were then transferred to a temperature and humidity-controlled incubator set to 21 ± 1°C. From 0-6 days post fertilization (6 dpf), water in all bowls was replaced 2-3 times a day. Beginning at 6 dpf, larvae were transferred from the incubator to the aquatics facility. Larvae were then fed twice a day with live *Artemia* (brine shrimp). At 9 dpf, larvae were tested for stress. As tracking in the transparent cavefish is challenging, 9 dpf was chosen because this was the youngest stage for which we could still track animals reliably. All methods and experiments in this study were approved by the Institutional Animal Care and Usage Committee (IACUC) at Florida Atlantic University (protocol number A17-21).

### Behavioral measures of stress using shock

The shock assay was carried out as previously described in zebrafish (Duboué et al., 2017), with minor modifications to use in *A. mexicanus* fry. A 6cm^3^ chamber was constructed by placing by placing custom-printed four-walled outer barrier (MakerBot Replicator 2) atop a 0.5 cm clear acrylic bottom. A pair of stainless-steel electrodes were placed on either side of the box, and the electrodes were attached to a square pulse electrical stimulator (Grass S48D, Natas Neurology, RI). An elevated platform was mounted in the center of the chamber, and a 40μm cell strainer (Corning, Durham, NC. Inner diameter=20mm, height of water=6mm) was positioned on the platform. The entire chamber was positioned on top of a custom designed infrared light source (880 nm), and a high speed charged couple device (CCD) camera (Grasshopper 3, Point Grey) fitted with a fixed focus lens (75 mm DG Series fixed focus lens, Edmond Optics) was positioned on top. The camera was connected to a computer (Dell XPS, core i7) and videos were recorded using FlyCap2 software (Point Grey) at 60 frames per second. To examine their response to shock, a single 9 dpf *A. mexicanus* fry was placed centrally in the cell strainer and allowed to acclimate for 1-min, and the video camera was activated. Fish were recorded for 1-min, and then 5 electric shocks were delivered (28V, 1 pulse per second, 200 msec pulse duration). Locomotor activity was then recorded for an additional minute. The protocol for assessing fish behavior were taken from those originally described for zebrafish (Duboué et al., 2017).

Videos were analyzed offline by first tracking individual larvae using Ethovision XT software (v13, Noldus). Tracking data was then inspected by a trained experimenter to ensure that tracking data accurately reflected the locomotor behavior of the larva, and when discrepancies were found, the tracks were modified to accurately reflect the (x,y) position of the fish using the ‘track editing’ function in Ethovision XT. All (x,y) pixel coordinates per frame were exported from Ethovision and imported into MATLAB (Mathworks). Using custom written scripts in MATLAB, we determined locomotor activity over time as well as total freezing. Total freezing duration was determined as any period of inactivity equal to or longer than 1.99 sec (Duboué et al., 2017).

### Measuring whole body cortisol levels

Whole body cortisol was measured using previously described protocols modified for *A. mexicanus* larvae (Cachat et al., 2010; Duboué et al., 2017). Briefly, 9 dpf larvae *A. mexicanus* were grouped in pools of 10 larvae per pool. Pools of larvae were transferred to 40μm cell strainer baskets (Corning, Durham, NC. Inner diameter=20mm, height of water=6mm). The baskets containing the larvae were transferred to the shock apparatus described above, and larvae were allowed 60 min to acclimate to the environment. After acclimation, larvae were shocked five times using the same Grass stimulator described above (28V, 1 pulse per second, 200 msec pulse duration). After stimulation, larvae were given 15 min recovery period, as this is the time that it takes for maximum activation of the cortisol pathway (Facchin et al., 2015). A control group of larvae were handled similarly, except no stimulation was delivered. Larvae were then euthanized using an overdose of MS222 (Sigma, cat. number A5040), and transferred to 1.5 mL Eppendorf tubes. Water was removed from the tubes, fish were flash frozen using liquid nitrogen, and the tubes were placed in a −20 °C freezer overnight. The following day, fish were homogenized in 1 mL of a 1X Phosphate Buffered Solution (PBS; Fisher, BP3994) using a pestle. Fifty μL of the homogenate was taken from each and stored for protein analysis (see below). The remaining homogenate was transferred to 4 mL glass scintillation vials. 2 mL of diethyl ether was added to the tube, and tubes were vortexed at high frequency for 1 min. Samples were then centrifuged at 5000 rpm at 4°C for 10 minutes. The top organic layer was removed using a glass pipette and transferred into a second 4 ml glass vial and kept under the fume hood. This process was repeated an additional two times. To expedite the rate at which diethyl ether evaporates, a small tube supplying nitrogen gas was added fitted into each tube, and nitrogen gas applied until the ether was evaporated. Following complete evaporation, the glucocorticoid homogenate was resuspended in 1 mL 1X PBS, and the vials were placed in a 4°C refrigerator overnight. On the following day, samples were assayed using a human salivary enzyme linked immunosorbent assay according (ELISA) to the manufacturer’s protocol (Salimetrics, cat. Number 1-3002, State College, PA). ELISA plates were then read on a spectrophotometer plate reader (FLOUstar Omega, BMG Labtech). The ELISA protocol was originally described for its use in zebrafish (Cachat et al., 2010), and we maintained those specifications including standards for *Astyanax*.

Whole body cortisol measures were normalized using total protein. Total protein was measured from the 50 μL sample of homogenate (described above) using a Quick Start Bradford assay (BioRad, Cat. Number 5000205). Bovine Serum Albumin (BSA) with concentrations 0.125 to 2.0mg/mL from a standard set served as protein standards (BioRad, Cat. Number 5000207). Then, 250μL of a 1X Bradford dye was added to 5 μL homogenate taken from each sample and mixed well in a 96 well plate (Corning, Cat. Number CLS3997). Samples were incubated for 5 min, and absorbance was determined using a spectrophotometer plate reader (FLOUstar Omega, BMG Labtech). All samples were converted from total absorbance to total protein concentration by relating absorbance to the BSA standards. Finally, total cortisol was normalized to total protein.

### Quantitative real-time polymerase chain reaction of crh, gr and mr

Expression analysis of *corticotropic releasing hormone*, the *glucocorticoid receptor* (*nuclear receptor subfamily 3 group C member 1*; *nr3c1*) and the *mineralocorticoid receptor* (*nuclear receptor subfamily 3 group C member 2*; *nr3c2*) were performed using quantitative real-time polymerase chain reaction (qRT-PCR). BLAST/BLAT (Ensembl) (Altschul et al., 1990) was first performed on the genes to ensure that there are no gene duplications and only a single gene isoform exist. Then, primers were designed for all three genes using Primer-BLAST (NCBI), and the regions amplified were ensured to be conserved in both surface and Pachon populations. Primer sequences were as follows: crh forward primer CCATCTCTCTGGACCTGACC; reverse primer: TCCATCATTCTGCGGTTGCT; gr forward primer: CGCCGAAATCATCAGCAACC; reverse primer: TAAGGCATGGTGTCCCGTTG; mr forward primer: TTCTTCAAGAGAGCGGTGGAA; reverse primer: GCCTGGAGACACTTCCGTA; beta-actin: GCTCTCTTCCAGCCTTCCTT; reverse: GCACTGTGTTGGCATACAGG. Larvae grown in glass bowls (Pyrex, 470mL) were sampled for analysis. RNA was extracted from pools of twenty larvae at 9 dpf with TRIzol (Fischer, cat number 15596018) and chloroform, and purified using the QIAGEN RNeasy mini kit. Then, RNA was reverse transcribed to cDNA using iScript Reverse Transcription Supermix (Bio-Rad, cat. number 1708840), and the cDNA was used in the qRT-PCR. The detection of fluorophore (SsoAdvanced Universal SYBR Green Supermix, cat number 172570, Bio-Rad) and thermal cycling were performed in the CFX96 Real-Time System (Bio-Rad) that is connected to the C1000 Touch Thermal Cycler (Bio-Rad). All genes analyzed were normalized to beta-actin.

### Statistics

All statistics were performed using Prism software (Graphpad, version 8) or Excel (Microsoft version 16). In the cases were pairwise comparisons of before shock and after shock were performed, a Wilcoxon paired t-test was used. In cases where more than 2 comparisons were examined, we used a Kruskal-Wallis test, and when differences were reported, this was followed by a Dunn’s test correcting for multiple comparisons. Correlational analysis was performed using a Spearman’s rank order correlation. For all statistical test, significance was determined as p being less than or equal to 0.05 alpha. All data will be made available upon request.

## Results

### Behavioral stress responses are reduced in Pachón cavefish

Electric shock has been used extensively in adult and larval zebrafish (Agetsuma et al., 2010; Valente et al., 2012; Tabor et al., 2014; Kenney et al., 2017). To determine whether larval cavefish have reduced responsiveness to stress compared to surface conspecifics, we assessed stress by applying mild electric shock and examining differences in freezing times pre- and post-stimulation. We subjected nine-day-old surface fish and compared their responses to shock to age matched Pachón fry. Briefly, larvae were placed individually in a chamber fitted with electrodes, and their locomotor activity recorded before and after electric stimulation (Fig. 1A). Similar to zebrafish, larval surface fish show an increase in freezing duration in the minute following stimulation (Fig. 1B, blue bars). By contrast, exposure to electric shock did not induce freezing in Pachón cavefish (Fig. 1B, black bars). These data reveal that reduced stress in *A. mexicanus* cavefish previously described in *A. mexicanus* adults, is also present in the larval stage.

**Figure 1.**
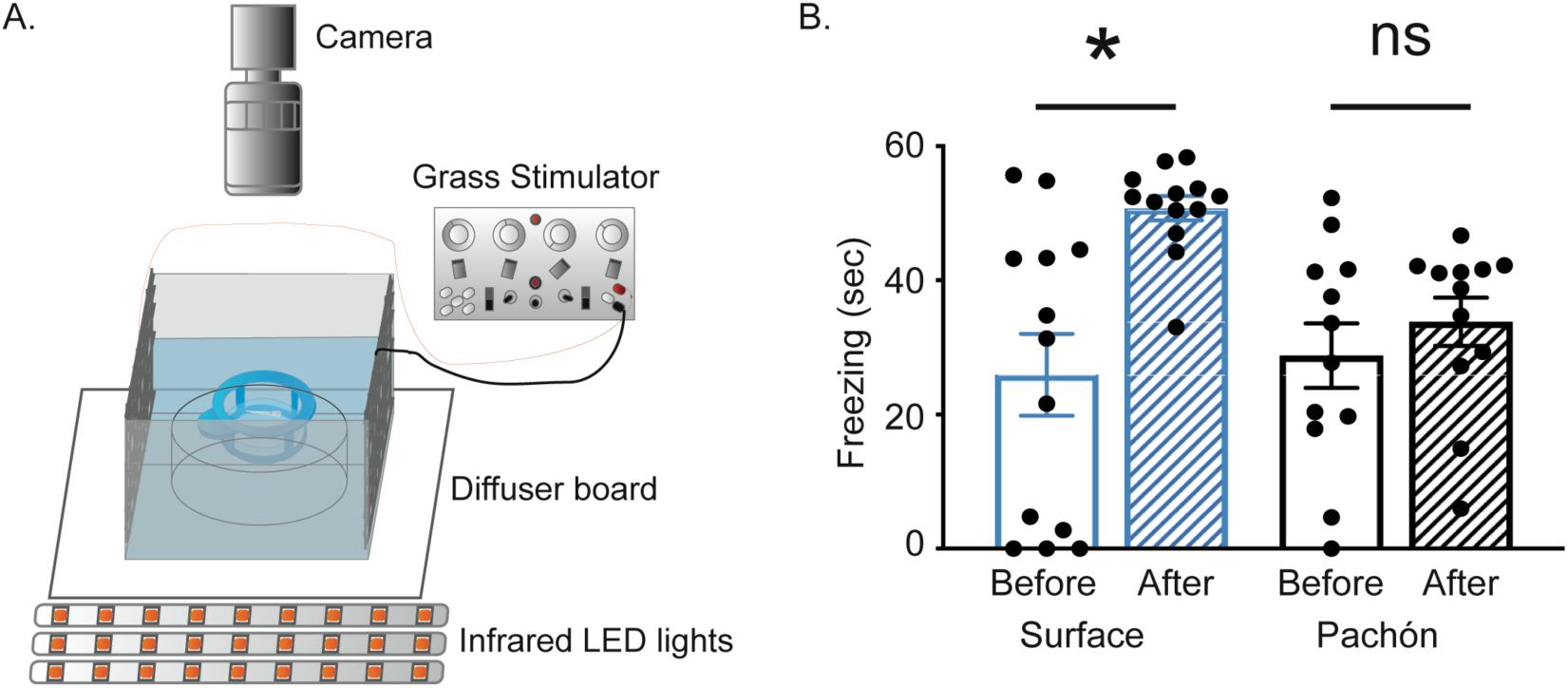
Pachón cavefish have dampened behavioral response to stress. (A) Diagram of shock assay. (B) Pachón cavefish freeze for less time relative to surface fish. Surface fish (blue) have increased freezing post-shock (hatched bar) relative to pre-shock freezing durations (solid bar) (Wilcoxon signed rank test, W=71.0, p=0.012). By contrast, freezing levels pre-(solid bars) and post-shock (hatched bars) were not significantly different for Pachón cavefish larvae (black bars) (Wilcoxon signed rank test, W=22.0, p=0.421). Error bars show ± standard error of the mean. * denote significance below p=0.05. ns denotes no significance.

### Diminished activation of the hypothalamic-pituitary-interrenal (HPI) axis in cavefish

A hallmark of stress is elevated levels of glucocorticoid stress hormones, such as cortisol in humans and fish, and corticosterone in rodents (Alsop and Vijayan, 2008, 2009). At the presentation of a stressful cue, hypothalamic neurons expressing *corticotropic releasing hormone* (*crh*) direct the interrenal gland that is embedded in the head kidney, to produce and release glucocorticoids into the blood stream (Collier et al., 2017), and glucocorticoids signal back to the brain by binding to glucocorticoid and mineralocorticoid receptors (Collier et al., 2017). To determine the response of the HPI axis to a stressful stimulus we measured cortisol levels in 9 dpf surface fish and Pachón cavefish larvae following exposure to shock. We found that, compared to unshocked controls, surface animals had a significant increase in cortisol 15-min following stimulation (Fig. 2A, blue bars). Conversely, Pachón cavefish had a small increase in cortisol, which was not significantly different than unshocked controls (Fig. 2A, black bars). Analysis of cortisol levels at 30-min or 1-hr post shock did not indicate a delayed response in Pachón cavefish, and in surface larvae, cortisol levels were comparable to before shock by 30-min (Supp. Fig. 1). These data demonstrate that, in addition to diminished behavioral responses to stress, Pachón cavefish have diminished physiological markers of stress as well.

**Figure 2.**
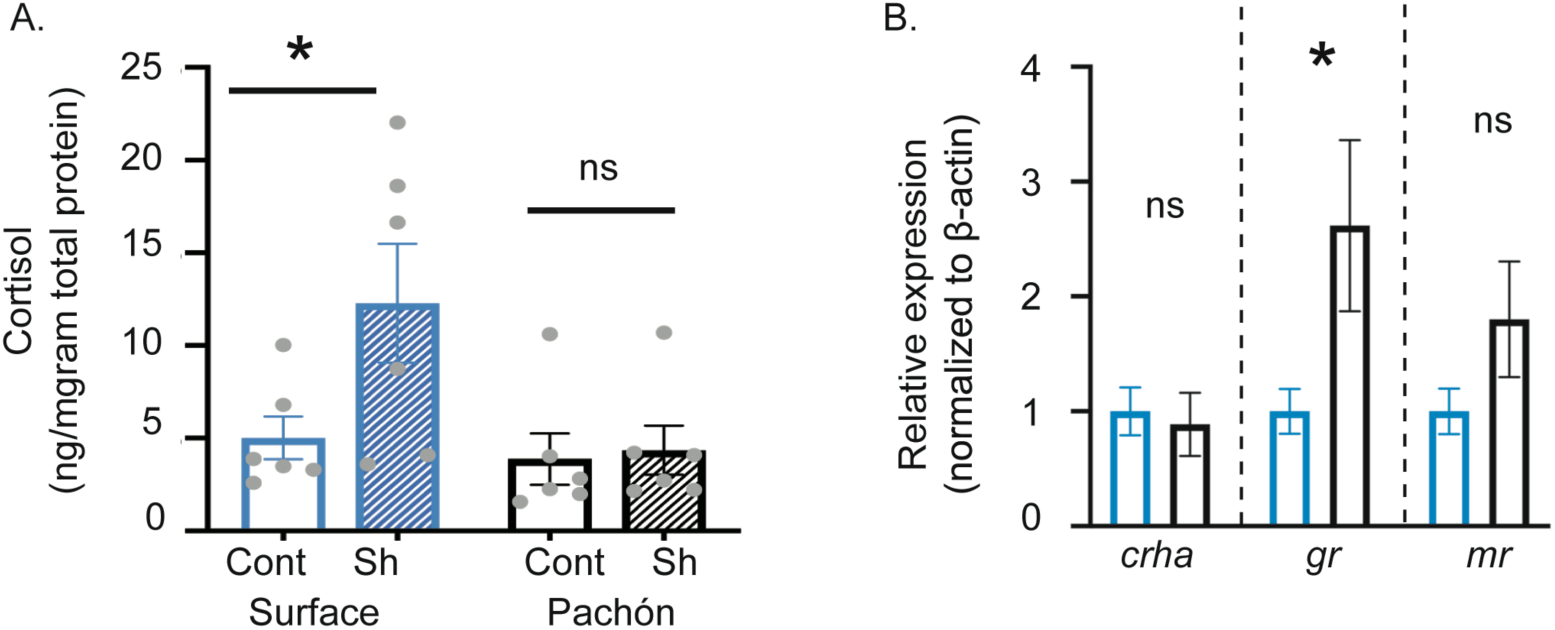
Pachón cavefish have reduced activity of the neuroendocrine stress axis. (A) In response to stress, surface larvae exhibit an increase in the stress hormone cortisol (Wilcoxon signed rank test, W=30.0, p=0.0078); however, in response to the same stimulus, Pachón larvae show no significant changes in whole body cortisol levels (Wilcoxon signed rank test, W=18.0, p=0.25). (B) Quantification of genes in the hypothalamic-pituitary-interrenal axis using qPCR. Measures of corticotropic releasing hormone (crha) were not significantly different between cave and surface fish (student’s t-test, t=0.5386, p=0.602). Expression of the glucocorticoid receptor (gr) was higher in cavefish relative to surface fish (student’s t-test, t=2.559, p=0.028). No significant differences were observed in the expression of the mineralocorticoid receptor (mr) (student’s t-test, t=1.201, p=0.257). For all plots, blue bars represent surface fish and black bars represent cavefish. ‘Cont’ represents controls and ‘Sh’ represents shocked larvae. Error bars show ± standard error of the mean. * denote significance below p=0.05. ns denotes no significance.

We next quantified differences in mRNA expression of genes in the HPI axis in surface and Pachón cavefish. We first examined the expression of *crha*, a gene that is upstream of cortisol and predominantly responsible for cortisol activation, and found no significant differences (Fig. 2B). We next examined the expression level of cortisol receptors. The mineralocorticoid receptor is a high-affinity receptor that is considered to be saturated with low levels of cortisol, and the low-affinity glucocorticoid receptor, which becomes activated upon high levels of cortisol (Reul and De Kloet, 1985; Reul et al., 1987). The glucocorticoid receptor is a nuclear receptor, which suppresses the HPI axis through a negative feedback loop mechanism. We found that expression of glucocorticoid receptor was significantly higher in Pachón cavefish than surface fish, but no significant differences in the mineralocorticoid receptor. Together, these data suggest elevated levels of glucocorticoid receptor suppress cortisol levels in cavefish.

### F2 analysis reveals independence of stress and morphological traits

The ability to breed surface/cave and cave/cave hybrids allows for examining how morphological and behavioral traits segregate. To explore whether the diminished stress response in cave animals is dominant or recessive, we crossed a pure-breeding surface fish to a pure-breeding Pachón fish to produce a brood of F_1_ progeny. F_1_ fish were tested in the shock assay. The mean total freezing duration in F_1_ animals was not significantly different than freezing pre-shock, though total times were more variable (Fig. 3A, purple bars), suggesting a dominant cave trait. Next, we incrossed F_1_ fish and examined the F_2_ progeny in the shock assay. Total pre- and post-shock freezing times were significantly different in F_2_ animals (Fig. 3A, magenta bars). Importantly, the range of freezing times in F_2_ animals was significantly greater than either of the pure-breeding stocks (surface σ^2^= 42.97;Pachón σ^2^ = 154.0; F2 σ^2^= 283.92), and the range of post-shock freezing times spanned the range of surface to cave animals. These data reveal that diminished stress in Pachón animals is genetically encoded, and the genes in cavefish are mostly dominant to those in surface conspecifics.

**Figure 3.**
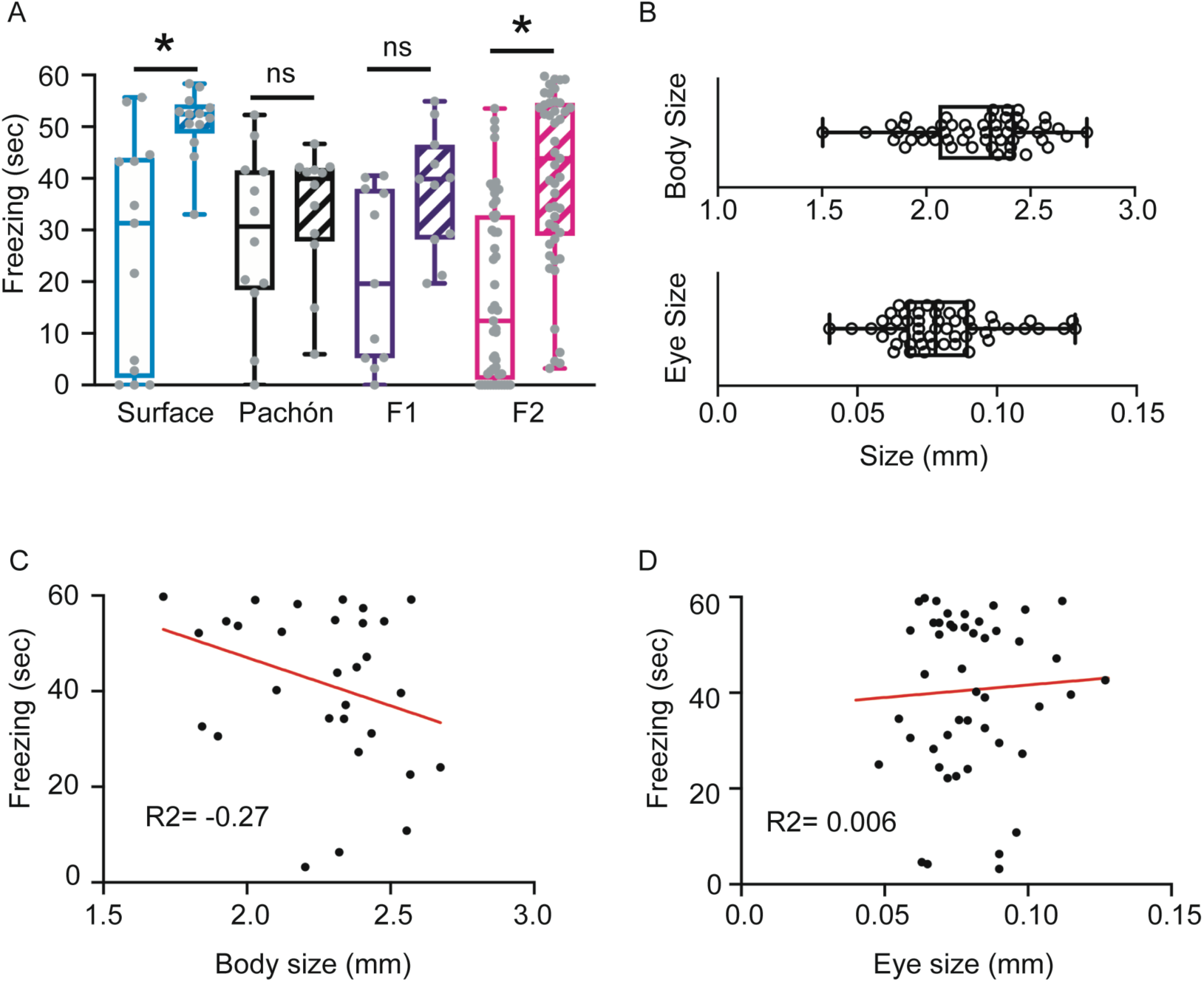
Reduced stress in cavefish is genetically determined, and does not segregate with body or eye size. (A) Distribution of freezing duration pre- and post-shock for surface fish (blue), Pachón cavefish (black), surface x cave F1 hybrids (purple) and surface x cave F2 hybrids (magenta). Freezing was significantly increased post shock for surface fish (Wilcoxon signed rank test, W=71.0, p=0.012), but not significantly different pre- and post-shock for Pachón cavefish (Wilcoxon signed rank test, W=22.0, p=0.421). Freezing durations pre- and post-shock for F1 hybrids approached significance (Wilcoxon signed rank test, W=42.0, p=0.067), and was intermediate between surface and cavefish. Freezing times pre- and post-shock for F2 hybrids was significant (Wilcoxon signed rank test, W=49.0, p<0.001), and spanned the range of surface and cave post-shock freezing. (B) Distribution of body length and eye size in surface x Pachón cave F2 hybrids. (C) The length of the body in F2 hybrids was not correlated significantly with freezing duration post-shock (Spearman’s test, rho=-0.27, p=0.14). (D) Eye size in F2 hybrids was not correlated significantly with freezing duration post-shock (Spearman’s test, rho=0.006, p=0.96). Error bars in A and B depict the max to min range. Red lines in C and D show the best fit in a linear regression analysis. * denote significance below p=0.05. ns denotes no significance.

To determine whether the stress responses are related to morphological phenotypes, for example, their ability to see, we tested the relationship between stress to eye size and body size in F_2_ hybrid fish. We first looked at both eye size, since visual cues could affect stress responses, and body size, as the size of the body could alter stimulus sensation. Consistent with previous reports, body size is smaller in age-matched surface animals compared to cavefish, whereas eye size is smaller in cavefish relative to surface. We measured both body and eye size in the F_2_ fish that we tested for stress, and found that these values were variable (Fig. 3B). We next asked whether stress was correlated with either morphological trait. A linear correlation of body size to freezing reveal a negative slope (m = −20.18), whereas a linear correlation of eye size to freezing revealed a positive slope (m = 52.29). However, a statistical correlation analysis revealed that neither body size nor eye size was significantly correlated with freezing times (body size∼ freezing: Spearman’s ρ^2^= −0.27, p>0.05; eye size ∼ freezing: Spearman’s ρ^2^ = 0.006, p>0.05), suggesting that reduced stress in cave animals is a novel trait, which is likely independent of these two morphological attributes.

### Convergence of stress in some, but not all larval A. mexicanus populations

The multiple independently evolved cavefish populations of *A. mexicanus* provide a unique opportunity to address whether evolution repeatedly has altered different traits, or whether the evolved differences are unique to some cavefish populations and not others. To assess whether or not cavefish from other localities also have dampened stress responses, we raised larvae from the Tinaja and Molino caves. Previous studies suggest that Tinaja are more closely related to fish from the Pachón cave and other populations from the El Abra region, whereas individuals from the Molino cave are evolved from a “newer” colonization of surface fish from the Sierra Guatemala region and are more divergent (Bradic et al., 2012). Surface fish subjected to shock exhibit a significant increase in freezing times, consistent with them having a characteristic stress response (Fig. 4, blue bars). When larvae derived from the Tinaja cave were tested, they showed little freezing post-shock (Fig. 4, pink bars), similar to our observations with Pachón larvae. By contrast, larvae from the Molino cave showed an increase in freezing similar to what we observed for surface fish (Fig. 4, orange bars). Interestingly, analysis of Molino cavefish behavior pre-stimulus revealed significantly diminished basal freezing or immobility compared to Tinaja; and also trended towards diminished stress compared to surface fish, although the differences were not significant (p=0.093). Conversely, Tinaja cavefish froze significantly less than Molino cavefish after shock, despite having higher basal freezing, suggesting a divergence in stress responses among different cavefish populations at larval stages. Taken together, these data suggest that dampened stress responses are conserved across developmental time points for Pachón and Tinaja populations, and demonstrate a divergence in these dampened responses in Molino, whose baseline activity is higher than both Tinaja and Pachón, and where an increase in freezing following shock is apparent in larval stages.

**Figure 4.**
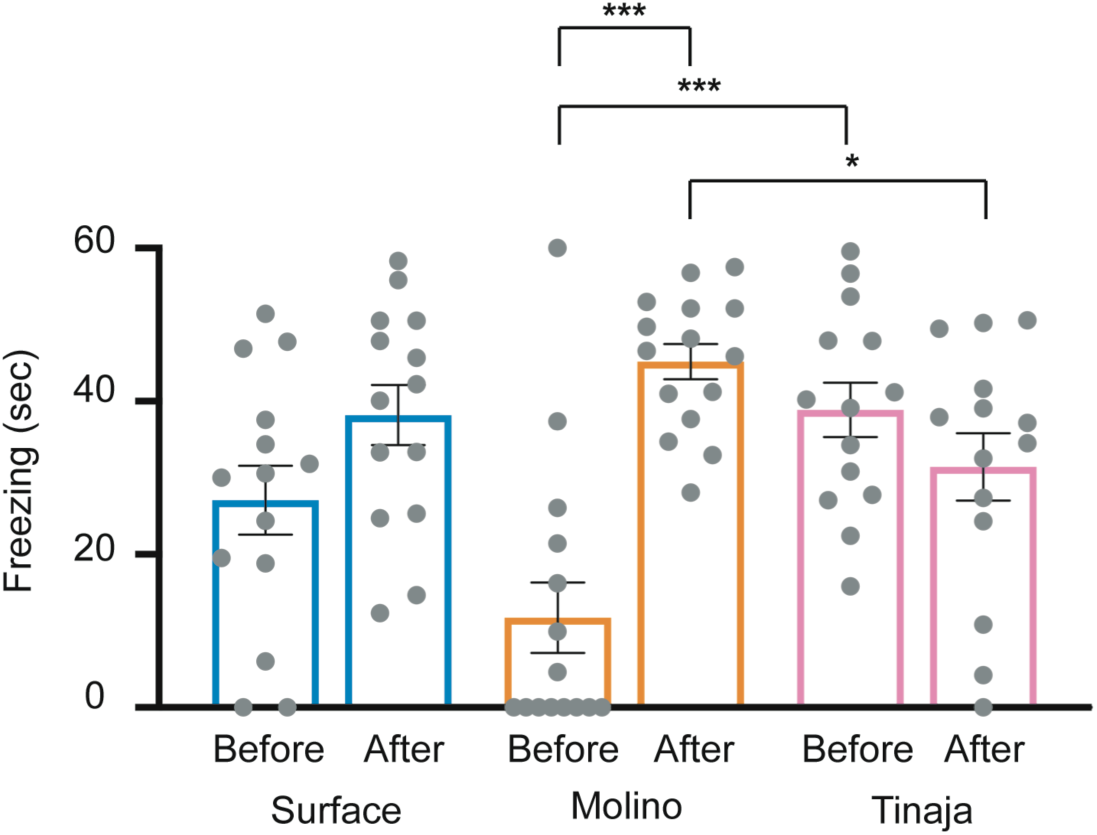
Divergence of stress responses in different, independently evolved populations of cavefish. Surface fish (blue bars) show more freezing post-shock compared to before stimulation (Wilcoxon signed rank test, W=62.0, p=0.049). Molino cavefish (orange bars) show significantly higher levels of freezing post-shock relative to pre-stimulation (Wilcoxon signed rank test, W=116.0, p=0.0002), whereas Tinaja cavefish (pink bars) show no significant differences (Wilcoxon signed rank test, W=-51.0, p=0.12). Comparing basal activity pre-stimulation, Molino cavefish show significantly reduced levels of freezing compared to Tinaja cavefish (Kruskal-Wallis test, p=0.0004), yet post-stimulus, Tinaja cavefish have reduced freezing compared to Molino cavefish (Kruskal-Wallist test, p=0.0423).

## Discussion

In this study, we have demonstrated that larval cavefish from some but not all caves have evolved to have a diminished stress response relative to their more ancestral surface fish populations. Moreover, we have found that Pachón cavefish have an elevated expression of the glucocorticoid receptor. These data not only corroborate previous studies demonstrating reduced stress in adult populations (Chin et al., 2018), but they also reveal a change in stress behaviors from larval to adult in Molino populations.

### Ontology of stress in Astyanax

In mammals, including rodents and humans, stress responses can vary considerably across development (Adriani and Laviola, 2004; Paus et al., 2008). Several studies, for example, have demonstrated that young or adolescent individuals have different degrees of stress from their adult counterparts (Slawecki, 2005; Hefner and Holmes, 2007; Paus et al., 2008; Lynn and Brown, 2010). Whereas the ontological reasons for these differences remains poorly understood, one prominent hypothesis is that the neuronal circuits and physiological mechanisms that modulate these behaviors are not fully developed at these early stages of development (McCormick and Mathews, 2007; Zimmermann et al., 2019). Our data demonstrate that some cavefish populations such as Pachón and Tinaja have diminished stress at both larval and adult stages, whereas others such as Molino have stress responses similar to surface fish as larvae, but then as adults have a diminished response. These data show clear ontological differences in stress between cavefish populations. For example, differing environmental demands of the Molino cave compared to the caves in which Pachón and Tinaja inhabit could account for these differences (Mitchell et al., 1977). Similarly, differences in genetic variation could also account for the ontological variations. Tinaja and Pachón are derived from an older stock of surface fish, whereas Molino are derived from a separate invasion from a newer stock of fish (Bradic et al., 2012). Moreover, many traits such as eye loss have arisen in new and old stocks through different genetic mechanisms (Borowsky, 2008b). How genetic differences between old- and new-invasion cavefish contribute to differences in the ontology of stress remain unknown, but the naturally occurring genetic variation in these stocks provides a powerful model to examine how genes can contribute to stress ontology.

The demonstration of stress in larval cavefish has considerable implications for establishing cavefish as a model for examining stress. The small size, simplified nervous system, and transparency, together with robust behavioral responses make larval fish especially attractive for evaluating how the brain drives behavior (Friedrich et al., 2010, 2013; Fero et al., 2011; Ahrens and Engert, 2015). Moreover, with transgenic technology being introduced recently into the *Astyanax* model (Stahl et al., 2019b; a), the system should be a powerful tool to assess how evolution has impacted the genes and development of neuronal circuits modulating stress in vertebrates.

### Glucocorticoid receptor mediates reduced stress

The molecular basis for reduced stress in cavefish animals remains unclear, yet our data demonstrating that glucocorticoid receptor is increased points to a role for the modulation of physiological mechanisms of stress. Across vertebrates, stress is modulated by a brain-to-periphery pathway. In mammals, stressful stimuli activate neurons in the hypothalamus that utilize corticotropic releasing hormone (crh). *crh* signals to the anterior pituitary, which then signals to the adrenal gland to produce and secrete glucocorticoids such as corticosterone or cortisol. Once produced, cortisol feeds back to the brain, and through binding to its glucocorticoid receptor (gr), it downregulates the activity of *crh* neurons. Our data reveal that cavefish have diminished stress behaviors, lower cortisol and a higher expression of the glucocorticoid receptor, suggesting that the increase in *gr* expression could, in part account for the diminished stress behaviors. It is not clear what is causing the increased expression of the glucocorticoid receptor, but future work should help to elucidate these mechanisms.

### Genetic contributions as demonstrated by F_1_ and F_2_ data, and the F_2_ correlation data

A powerful attribute to examining behavioral evolution in blind Mexican cavefish is the ability to cross breed fish from different populations, including the generation of surface x cave hybrids as well as crosses of cavefish from different cave localities. This ability allows researchers to examine pleiotropy through co-segregation with other traits, as well as an ability to perform quantitative genetic analyses (Protas et al., 2008; Yoshizawa et al., 2015; Jaggard et al., 2017). For example, differences in stress could be due, partly, to the differences in surface and cavefish’s ability to see. Alternatively, altered stress could correlate with differences in metabolism or size of the animal. Our data demonstrate an intermediate stress phenotype in surface cave F_1_ hybrids, as well as a distribution of stress responses in F_2_ hybrids that spans the range of surface to cave. These data not only point to a strong genetic underpinning of the differences in stress seen between surface and cave animals, but also suggest, given the F_1_ data, that cave alleles exhibit either intermediate dominance or a complex relationship with surface alleles.

In human populations, stress and other anxiety related disorders, are caused by both genetic and environmental factors (Dick, 2011; Sharma et al., 2016). Several genome wide association studies (GWAS) have been conducted in search of genetic variants that predispose patients to anxiety and stress related disorders, and the heritability of these disorders has been calculated to be between 30-50% (True et al., 1993; Stein et al., 2002; Sartor et al., 2012). However, it remains unclear how standing genetic variation contributes to the differences in susceptibility of anxiety and stress related disorders. Because the cavefish are a rare model where standing genetic variation is largely unimpeded, this model could be powerful in establishing how genetics lead to susceptibility of these and other psychological diseases.

### Neuroanatomical regions that modulate stress may be different between surface and cave morphs

Stress is modulated by myriad neuroanatomical loci, including the hypothalamus, the amygdala, hippocampus, preoptic area of the hypothalamus, and habenulae (Tovote et al., 2015), and many of these regions have been shown to differ significantly between cave and surface morphs in neural activity and/or neuroanatomy (Menuet et al., 2007; Alié et al., 2018; Jaggard et al., 2018; Loomis et al., 2019). For example, in fish and mammals, the hypothalamus has a critical role in the modulation and induction of stress (Spiess et al., 1981; Callahan et al., 1992; Cachat et al., 2010; Steenbergen et al., 2012; Wamsteeker Cusulin et al., 2013; Yeh et al., 2013). A recently published neuroanatomical brain atlas in cavefish shows that various subnuclei within the hypothalamus are morphometrically different (Loomis et al., 2019). Similarly, these data have revealed that the forebrain, an area which contains cell populations that are similar to the mammalian amygdala and hippocampus, is significantly expanded in cavefish populations (Alié et al., 2018; Loomis et al., 2019). These data suggest that areas that are essential to the proper modulation of stress are evolutionarily different between cave and epigean *Astyanax*.

Whereas the molecular mechanisms underlying these differences in neuroanatomy are not fully understood, several studies have pointed to key developmental factors that may play a role. For example, the genes *NK homeobox 2*.*1a* (*Nkx2*.*1a*) *LIM homeobox protein 6* (*lhx6*) has an important role in the development of the forebrain (Alié et al., 2018). In cavefish, the expression of both *nkx2*.*1a* and *lhx6* are significantly expanded as compared to surface fish, suggesting that these factors may be involved in the evolutionary modification of the telencephalon. Similarly, the hypothalamus is modulated by numerous molecular factors, but one that is particular important is *sonic hedgehog* (*shh*) (Corman et al., 2018). Expression of *shh* has been shown to be expanded both along the midline as well is in the hypothalamus (Menuet et al., 2007). Expanded expression of this gene may be important for the resistance to stress that we report in cavefish. Together, these data suggest that modifications in the molecular mechanisms underlying brain development could be important in evolutionarily derived behaviors.

### Functional implications for diminished stress in cavefish

Many caves, including some in the Sierra de el Abra region of Mexico where cavefish are found, are characterized by perpetual darkness, a near absence of primary producer, low food availability, and a reduction of predators (Poulson and White, 1969; Mitchell et al., 1977; Culver, 1982). In the caves of Northeast Mexico, cave *Astyanax* are thought to be at the top of their food chain, and to have little to no predatory threat. Predatory threat has been shown to correlate significantly with stress responses (Clinchy et al., 2013). For example, in both Belding ground squirrels as well as the mosquito fish, *Gambusia hubbsi*, animals raised in a predator rich environment have been shown to have increased stress responses, compared to siblings that were raised with low predator threat (Mateo, 2007; Heinen-Kay et al., 2016). In Trinidadian guppies, increased predator threat promotes social relationships, and predator stimuli over time induced population changes towards ‘shy’ behaviors such as increased freezing (Heathcote et al., 2017; Houslay et al., 2018). These data reveal that even in unrelated species, predator threat can cause differences in stress. Our data suggest that both adult (Chin et al., 2018) and some populations of larval fish follow this trend. Moreover, our data reveal that predator threat can not only correlate with differences in stress, but also show that predator threat may be a powerful evolutionary driving force. We hypothesize that the low predator threat of the *Astyanax* caves either resulted in relaxed selection of stress responses, or may be a selective driving force itself. Further experiments will reveal whether this is the case, and if so, whether or not diminished stress in cavefish is selected for, or emerged through neutral drift.

### Future directions

The blind Mexican cavefish has emerged in recent years as a powerful system to examine how naturally occurring genetic variation contributes to various diseases such as obesity (Aspiras et al., 2015), diabetes (Riddle et al., 2018), insomnia (Duboué et al., 2011; Jaggard et al., 2018) and heart regeneration (Stockdale et al., 2018). Our data suggest that the cavefish may also be a powerful system to examine how naturally occurring genetic variation contributes to differences in stress, and could be important for determining genetic predispositions to stress and anxiety related disorders. In addition, understanding stress and its effects in the long term on growth, cardiovasculature, metabolism, and musculature is important in aquaculture (Sadoul and Vijayan, 2016). We have previously demonstrated that adult *A. mexicanus* have diminished stress responses (Chin et al., 2018), and these data extend this to larval forms. We expect that future work using powerful approaches unique to larvae, such as whole brain functional imaging and tracing of neural circuits that modulate stress, together with genotype-phenotype associations between genes and stress responses will shed significant light into genetic pre-disposition to stress disorders.

## Figures

**Supplemental Figure 1.**
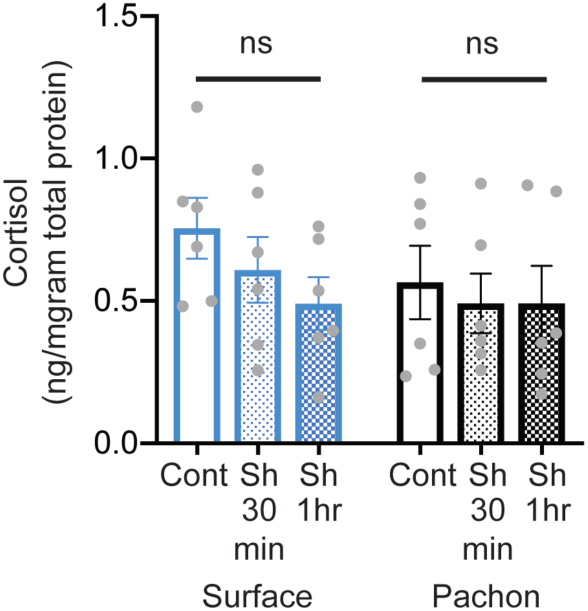
Stress responses are diminished by 30 minutes in surface fish larvae, and no delayed responses are observed in Pachón cavefish. Cortisol measurements done at 30-min and 1-hr post-shock were not significantly different than control siblings in both populations (Wilcoxon signed rank test, Surface-Cont vs. Sh 30min: W=-7.0, p=0.563. Cont vs. Sh 1hr: W=-9.0, p=0.438. Pachón-Cont vs. Sh 30min: W=-7.0, p=0.563. Cont vs. Sh 1hr: W=-9.0, p=0.438.). ‘Cont’ represents controls and ‘Sh’ represents shocked larvae. Blue bars represent surface fish and black bars represent Pachón cavefish. Error bars show ± standard error of the mean.

## References

Adriani W, Laviola G. 2004. Windows of vulnerability to psychopathology and therapeutic strategy in the adolescent rodent model. Behav Pharmacol 15:341–52.

Agetsuma M, Aizawa H, Aoki T, Nakayama R, Takahoko M, Goto M, Sassa T, Amo R, Shiraki T, Kawakami K, Hosoya T, Higashijima S, Okamoto H. 2010. The habenula is crucial for experience-dependent modification of fear responses in zebrafish. Nat Neurosci 13:1354–1356.

Ahrens MB, Engert F. 2015. Large-scale imaging in small brains. Curr Opin Neurobiol 32:78–86.

Alié A, Devos L, Torres-Paz J, Prunier L, Boulet F, Blin M, Elipot Y, Retaux S, Alie A, Devos L, Torres-Pas J, Prunier L, Boulet F, Blin M, Elipot Y, Reteaux S. 2018. Developmental evolution of the forebrain in cavefish, from natural variations in neuropeptides to behavior. Elife 7. pii: e3.

Alsop D, Vijayan M. 2009. The zebrafish stress axis: Molecular fallout from the teleost-specific genome duplication event. Gen Comp Endocrinol 161:62–6.

Alsop D, Vijayan MM. 2008. Development of the corticosteroid stress axis and receptor expression in zebrafish. Am J Physiol - Regul Integr Comp Physiol 294:R711–9.

Altschul SF, Gish W, Miller W, Myers EW, Lipman DJ. 1990. Basic local alignment search tool. J Mol Biol 215:403–10.

Aspiras A, Rohner N, Marineau B, Borowsky R, Tabin J. 2015. Melanocortin 4 receptor mutations contribute to the adaptation of cavefish to nutrient-poor conditions. Proc Natl Acad Sci 112:9688–73.

Biran J, Tahor M, Wircer E, Levkowitz G. 2015. Role of developmental factors in hypothalamic function. Front Neuroanat 9:47:doi: 10.3389/fnana.2015.00047.

Borowsky R. 2008a. Astyanax mexicanus, the blind Mexican cave fish: A model for studies in development and Morphology. Cold Spring Harb Protoc 3: 11:doi:10.1101/pdb.emo107.

Borowsky R. 2008b. Restoring sight in blind cavefish. Curr Biol 18:R23–4.

Borowsky R. 2008c. Breeding Astyanax mexicanus through natural spawning. Cold Spring Harb Protoc:pdb.prot5091.

Bradic M, Beerli P, García-De Leán FJ, Esquivel-Bobadilla S, Borowsky RL. 2012. Gene flow and population structure in the Mexican blind cavefish complex (Astyanax mexicanus). BMC Evol Biol 12:9:doi: 10.1186/1471-2148-12-9.

Bradic M, Teotónio H, Borowsky RL. 2013. The population genomics of repeated evolution in the blind cavefish astyanax mexicanus. Mol Biol Evol 30:2383–2400.

Cachat J, Stewart A, Grossman L, Gaikwad S, Kadri F, Chung KM, Wu N, Wong K, Roy S, Suciu C, Goodspeed J, Elegante M, Bartels B, Elkhayat S, Tien D, Tan J, Denmark A, Gilder T, Kyzar E, DiLeo J, Frank K, Chang K, Utterback E, Hart P, Kalueff A V. 2010. Measuring behavioral and endocrine responses to novelty stress in adult zebrafish. Nat Protoc 5:1786–1799.

Callahan MF, Thore CR, Sundberg DK, Gruber KA, O’Steen K, Morris M. 1992. Excitotoxin paraventricular nucleus lesions: stress and endocrine reactivity and oxytocin mRNA levels. Brain Res 597:8–15.

Campos AC, Fogaça M V., Aguiar DC, Guimarães FS. 2013. Animal models of anxiety disorders and stress. Rev Bras Psiquiatr 35:S101–11.

Chin JS, Gassant CE, Amaral PM, Lloyd E, Stahl BA, Jaggard JB, Keene AC, Duboue ER. 2018. Convergence on reduced stress behavior in the Mexican blind cavefish. Dev Biol 441:319–327.

Clinchy M, Sheriff MJ, Zanette LY. 2013. Predator-induced stress and the ecology of fear. Funct Ecol 27:56–65.

Cockrem JF. 2013. Individual variation in glucocorticoid stress responses in animals. Gen Comp Endocrinol 181:45–58.

Collier AD, Kalueff A V., Echevarria DJ. 2017. Zebrafish Models of Anxiety-Like Behaviors. In: Kalueff A V., editor. The rights and wrongs of zebrafish: Behavioral phenotyping of zebrafish. 1st ed. Springer International Publishing.

Corman TS, Bergendahl SE, Epstein DJ. 2018. Distinct temporal requirements for Sonic hedgehog signaling in development of the tuberal hypothalamus. Development 145:pii: dev167379.

Cornelis MC, Nugent NR. 2010. Genetics of Post-Traumatic Stress Disorder: Review and Recommendations for Genome-Wide Association Studies. Curr Psychiatr Rep 12:313–326.

Culver D. 1982. Cave life: evolution and ecology. 1st ed. Harvard University Press.

Dick DM. 2011. Gene-Environment Interaction in Psychological Traits and Disorders. Annu Rev Clin Psychol 7:383–409.

Dowling TE, Martasian DP, Jeffery WR. 2002. Evidence for multiple genetic forms with similar eyeless phenotypes in the blind cavefish, Astyanax mexicanus. Mol Biol Evol 19:446–55.

Duboué ER, Hong E, Eldred KC, Halpern ME. 2017. Left Habenular Activity Attenuates Fear Responses in Larval Zebrafish. Curr Biol 27:2154-2162.e3.

Duboué ER, Keene AC, Borowsky RL. 2011. Evolutionary convergence on sleep loss in cavefish populations. Curr Biol 21:671–676.

Elipot Y, Hinaux H, Callebert J, Rétaux S. 2013. Evolutionary shift from fighting to foraging in blind cavefish through changes in the serotonin network. Curr Biol 23:1–10.

Etkin A, Wager TD. 2007. Functional neuroimaging of anxiety: a meta-analysis of emotional processing in PTSD, social anxiety disorder, and specific phobia. Am J Psychiatry 164:1476–1488.

Facchin L, Duboué ER, Halpern MEE, Duboue ER, Halpern MEE. 2015. Disruption of Epithalamic Left-Right Asymmetry Increases Anxiety in Zebrafish. J Neurosci 35:15847–15859.

Fero K, Yokogawa T, Burgess HA. 2011. The behavioral repertoire of larval zebrafish. Neuromethods: 249–291.

Friedrich RW, Genoud C, Wanner AA. 2013. Analyzing the structure and function of neuronal circuits in zebrafish. Front Neural Circuits 7:doi: 10.3389/fncir.2013.00071.

Friedrich RW, Jacobson GA, Zhu P. 2010. Circuit Neuroscience in Zebrafish. Curr Biol 20:R371–81.

Gross JB. 2016. Convergence and parallelism in Astyanax cave-dwelling fish. In: Pontarotti P, editor. Evolutionary Biology: Convergent Evolution, Evolution of Complex Traits, Concepts and Methods. Cham: Springer. p 105–119.

Hadany L, Beker T, Eshel I, Feldman MW. 2006. Why is stress so deadly? An evolutionary perspective. Proc R Soc B Biol Sci 273:881–5.

Heathcote RJP, Darden SK, Franks DW, Ramnarine IW, Croft DP. 2017. Fear of predation drives stable and differentiated social relationships in guppies. Sci Rep 7:doi: 10.1038/srep41679.

Hefner K, Holmes A. 2007. Ontogeny of fear-, anxiety- and depression-related behavior across adolescence in C57BL/6J mice. Behav Brain Res 176:210–215.

Heinen-Kay JL, Schmidt DA, Stafford AT, Costa MT, Peterson MN, Kern EMA, Langerhans RB. 2016. Predicting multifarious behavioural divergence in the wild. Anim Behav 121:3–10.

Houslay TM, Vierbuchen M, Grimmer AJ, Young AJ, Wilson AJ. 2018. Testing the stability of behavioural coping style across stress contexts in the Trinidadian guppy. Funct Ecol 32:424–438.

Jaggard JB, Stahl BA, Lloyd E, Prober DA, Duboue ER, Keene AC. 2018. Hypocretin underlies the evolution of sleep loss in the Mexican cavefish. Elife 7:pii: e32637.

Jaggard JJ, Robinson BBG, Stahl BAB, Oh I, Masek P, Yoshizawa M, Keene ACA. 2017. The lateral line confers evolutionarily derived sleep loss in the Mexican cavefish. J Exp Biol 220:284–293.

Jeffery WR. 2009. Regressive Evolution in Astyanax Cavefish. Annu Rev Genet 141:520–529.

Keene A, Yoshizawa M, McGaugh S. 2015. Biology and Evolution of the Mexican Cavefish. 1st Editio. New York: Academic Press.

Kenney JW, Scott IC, Josselyn SA, Frankland PW. 2017. Contextual fear conditioning in zebrafish. Learn Mem 24:516–523.

Koenen KC, Nugent NR, Amstadter AB. 2008. Gene-environment interaction in posttraumatic stress disorder: Review, strategy and new directions for future research. Eur Arch Psychiatry Clin Neurosci 258:82–96.

Kowalko J, Rohner N, Linden T, Rompani S, Warren W, Borowsky R, Tabin C, Jeffery W, Yoshizawa M. 2013a. Convergence in feeding posture occurs through different genetic loci in independently evolved cave populations of Astyanax mexicanus. Proc Natl Acad Sci 110:16933–16938.

Kowalko JE, Rohner N, Rompani SB, Peterson BK, Linden TA, Yoshizawa M, Kay EH, Weber J, Hoekstra HE, Jeffery WR, Borowsky R, Tabin CJ. 2013b. Loss of schooling behavior in cavefish through sight-dependent and sight-independent mechanisms. Curr Biol 23:1874–1883.

Laberge F, Yin-Liao I, Bernier NJ. 2019. Temporal profiles of cortisol accumulation and clearance support scale cortisol content as an indicator of chronic stress in fish. Conserv Physiol 7:coz052.

Lacey JI, Lacey BC. 1958. Verification and extension of the principle of autonomic response-stereotypy. Am J Psychol 71:50–73.

Loomis C, Peuß R, Jaggard J, Wang Y, McKinney S, Raftopoulos S, Raftopoulos A, Whu D, Green M, McGaugh SE, Rohner N, Keene AC, Duboue ER. 2019. An adult brain atlas reveals broad neuroanatomical changes in independently evolved populations of Mexican cavefish. Front Neuroanat 13:doi: 10.3389/fnana.2019.00088.

Lynn DA, Brown GR. 2010. The ontogeny of anxiety-like behavior in rats from adolescence to adulthood. Dev Psychobiol 52:731–9.

Mateo JM. 2007. Ecological and hormonal correlates of antipredator behavior in adult Belding’s ground squirrels (Spermophilus beldingi). Behav Ecol Sociobiol 62:37–49.

Maximino C, de Brito TM, da Silva Batista AW, Herculano AM, Morato S, Gouveia A. 2010. Measuring anxiety in zebrafish: A critical review. Behav Brain Res 214:157–171.

Maximino C, Da Silva AWB, Araujo J, Lima MG, Miranda V, Puty B, Benzecry R, Picanco-Diniz DLW, Gouveia A, Oliveira KRM, Herculano AM. 2014. Fingerprinting of psychoactive drugs in zebrafish anxiety-like behaviors. PLoS One 9:e103943.

McCormick CM, Mathews IZ. 2007. HPA function in adolescence: Role of sex hormones in its regulation and the enduring consequences of exposure to stressors. Pharmacol Biochem Behav 86:220–33.

Menuet A, Alunni A, Joly J-SJ-S, Jeffery WR, Retaux S, Rétaux S. 2007. Expanded expression of Sonic Hedgehog in Astyanax cavefish: multiple consequences on forebrain development and evolution. Development 134:845–855.

Mitchell RWW, Russell WHH, Elliott WR, Elliot WR. 1977. Mexican Eyeless Characin Fishes, Genus Astyanax: Environment, Distribution, and Evolution. 1st ed. Lubbock, TX: Texas Tech Press.

Mueller T, Dong Z, Berberoglu MA, Guo S. 2011. The dorsal pallium in zebrafish, Danio rerio (Cyprinidae, Teleostei). Brain Res 1381:95–105.

Ornelas-García CP, Domínguez-Domínguez O, Doadrio I. 2008. Evolutionary history of the fish genus astyanax baird & Girard (1854) (Actinopterygii, Characidae) in mesoamerica reveals multiple morphological homoplasies. BMC Evol Biol 8:340.

Patton P, Windsor S, Coombs S. 2010. Active wall following by Mexican blind cavefish (Astyanax mexicanus). J Comp Physiol A Neuroethol Sensory, Neural, Behav Physiol 196:853–867.

Paus T, Keshavan M, Giedd JN. 2008. Why do many psychiatric disorders emerge during adolescence? Nat Rev Neurosci 9:947–57.

Poulson TL, White WB. 1969. The cave environment. Science (80-) 165:971–981.

Protas M, Tabansky I, Conrad M, Gross JB, Vidal O, Tabin CJ, Borowsky R. 2008. Multi-trait evolution in a cave fish, Astyanax mexicanus. Evol Dev 10:196–209.

Protas ME, Hersey C, Kochanek D, Zhou Y, Wilkens H, Jeffery WR, Zon LI, Borowsky R, Tabin CJ. 2006. Genetic analysis of cavefish reveals molecular convergence in the evolution of albinism. Nat Genet 38:107–111.

Reul JMHM, Van Den Bosch FR, De Kloet ER. 1987. Relative occupation of type-I and type-II corticosteroid receptors in rat brain following stress and dexamethasone treatment: Functional implications. J Endocrinol 115:459–67.

Reul JMHM, De Kloet ER. 1985. Two receptor systems for corticosterone in rat brain: Microdistribution and differential occupation. Endocrinology 117:2505–11.

Riddle MR, Aspiras AC, Gaudenz K, Peuß R, Sung JY, Martineau B, Peavey M, Box AC, Tabin JA, McGaugh S, Borowsky R, Tabin CJ, Rohner N. 2018. Insulin resistance in cavefish as an adaptation to a nutrient-limited environment. Nature 555:647–651.

Rohner N. 2018. Cavefish as an evolutionary mutant model system for human disease. Dev Biol 441:355–357.

Sadoul B, Vijayan M. 2016. Stress and Growth. In: Schreck C, Brauner C, editors. Biology of Stress in Fish. 1st ed. Academic Press. p 167–205.

Sakamoto T, Yoshiki M, Sakamoto H. 2017. The mineralocorticoid receptor knockout in medaka is further validated by glucocorticoid receptor compensation. Sci Data 4:170189:doi: 10.1038/sdata.2017.189.

Sartor CE, Grant JD, Lynskey MT, McCutcheon V V., Waldron M, Statham DJ, Bucholz KK, Madden PAF, Heath AC, Martin NG, Nelson EC. 2012. Common heritable contributions to low-risk trauma, high-risk trauma, posttraumatic stress disorder, and major depression. Arch Gen Psychiatry 69:293–9.

Schneiderman N, Ironson G, Siegel SD. 2005. Stress and Helth: Psychological, Behavioral, and Biological Determinants. Annu Rev Clin Psychol 1:607–628.

Schreck C, Tort L, Farrell A, Brauner C. 2016. Biology of Stress in Fish. 1st ed. Elsevier.

Sharma S, Powers A, Bradley B, Ressler KJ. 2016. Gene × Environment Determinants of Stress- and Anxiety-Related Disorders. Annu Rev Psychol 67:239–61.

Slawecki CJ. 2005. Comparison of anxiety-like behavior in adolescent and adult Sprague-Dawley rats. Behav Neurosci 119:1477–83.

Sneddon L, Wolfenden D, Thomson JS. 2016. Stress Management and Welfare. In: Schreck CB, Farrell LTA, Brauner C, editors. Biology of Stress in Fishes. 1st ed. Academic Press. p 167–205.

Spiess J, Rivier J, Rivier C, Vale W. 1981. Primary structure of corticotropin-releasing factor from ovine hypothalamus. Proc Natl Acad Sci U S A 78:6517–2.

Stahl BA, Jaggard JB, Chin JSR, Kowalko JE, Keene AC, Duboué ER. 2019a. Manipulation of Gene Function in Mexican Cavefish. J Vis Exp 146:doi: 10.3791/59093.

Stahl BA, Peuß R, McDole B, Kenzior A, Jaggard JB, Gaudenz K, Krishnan J, McGaugh SE, Duboue ER, Keene AC, Rohner N. 2019b. Stable transgenesis in Astyanax mexicanus using the Tol2 transposase system. Dev Dyn 248:679–687.

Steenbergen PJ, Metz JR, Flik G, Richardson MK, Champagne DL. 2012. Methods to quantify basal and stress-induced cortisol response in larval zebrafish. Zebrafish Protoc Neurobehav Res 66:121–141.

Stein MB, Jang KL, Taylor S, Vernon PA, Livesley WJ. 2002. Genetic and environmental influences on trauma exposure and posttraumatic stress disorder symptoms: A twin study. Am J Psychiatry 152:1675–81.

Stockdale WT, Lemieux ME, Killen AC, Zhao J, Hu Z, Riepsaame J, Hamilton N, Kudoh T, Riley PR, van Aerle R, Yamamoto Y, Mommersteeg MTM. 2018. Heart Regeneration in the Mexican Cavefish. Cell Rep 25:1997–2007.

Tabor KM, Bergeron SA, Horstick EJ, Jordan DC, Aho V, Porkka-Heiskanen T, Haspel G, Burgess HA. 2014. Direct activation of the Mauthner cell by electric field pulses drives ultrarapid escape responses. J Neurophysiol 112:834–44.

Tovote P, Fadok JP, Lüthi A. 2015. Neuronal circuits for fear and anxiety. Nat Rev Neurosci 16:317–331.

True WR, Rice J, Eisen SA, Heath AC, Goldberg J, Lyons MJ, Nowak J. 1993. A Twin Study of Genetic and Environmental Contributions to Liability for Posttraumatic Stress Symptoms. Arch Gen Psychiatry 50:257–64.

Valente A, Huang KH, Portugues R, Engert F. 2012. Ontogeny of classical and operant learning behaviors in zebrafish. Learn Mem 19:170–177.

Varga M, Ralbovszki D, Balogh E, Hamar R, Keszthelyi M, Tory K. 2018. Zebrafish Models of Rare Hereditary Pediatric Diseases. Diseases 6:pii: E43.

Wamsteeker Cusulin JI, Füzesi T, Watts AG, Bains JS. 2013. Characterization of Corticotropin-Releasing Hormone neurons in the Paraventricular Nucleus of the Hypothalamus of Crh-IRES-Cre Mutant Mice. PLoS One 8:e64943.

Wilkens H. 1971. Genetic Interpretation of Regressive Evolutionary Processes: Studies on Hybrid Eyes of Two Astyanax Cave Populations (Characidae, Pisces). Evolution (N Y).

Yeh CM, Glöck M, Ryu S. 2013. An optimized whole-body cortisol quantification method for assessing stress levels in larval zebrafish. PLoS One 8:e79406.

Yoshizawa M, Gorički Š, Soares D, Jeffery WR. 2010. Evolution of a behavioral shift mediated by superficial neuromasts helps cavefish find food in darkness. Curr Biol 20:1631–1636.

Yoshizawa M, Robinson BG, Duboué ER, Masek P, Jaggard JB, O’Quin KE, Borowsky RL, Jeffery WR, Keene AC. 2015. Distinct genetic architecture underlies the emergence of sleep loss and prey-seeking behavior in the Mexican cavefish. BMC Biol 13:doi: 10.1186/s12915-015-0119-3.

Zimmermann KS, Richardson R, Baker KD. 2019. Maturational changes in prefrontal and amygdala circuits in adolescence: implications for understanding fear inhibition during a vulnerable period of development. Brain Sci 9:pii: E65.

